# Unbiased method for spectral analysis of cells with great diversity of autofluorescence spectra

**DOI:** 10.1101/2023.07.28.550943

**Authors:** Janna E.G. Roet, Aleksandra M. Mikula, Michael de Kok, Cora H. Chadick, Juan J. Garcia Vallejo, Henk P. Roest, Luc J.W. van der Laan, Charlotte M. de Winde, Reina E. Mebius

## Abstract

Autofluorescence is an intrinsic feature of cells, caused by the natural emission of light by its cellular content, that can complicate analysis of flow cytometry data. Different cell types have different autofluorescence spectra and even within one cell type heterogeneity of autofluorescence spectra can be present, for example as a consequence of activation status or metabolic changes. By using full spectrum flow cytometry, the emission spectrum of a fluorochrome is captured by a set of detectors across a range of wavelengths, creating an unique spectrum for this fluorochrome, that is used to unmix the signal of a full stained sample into the signals of the different fluorochromes. Importantly, this technology can also be used to identify the autofluorescence signal of an unstained sample, which can be used for unmixing purposes and to separate the autofluorescence signal from the fluorophore signals. However, this only works if the sample has one homogeneous autofluorescence spectrum. To analyze samples with a heterogeneous autofluorescence spectral profile, we here setup an unbiased workflow to detect all different autofluorescence spectra present in a sample to take them along as ‘autofluorescence signatures’ during the unmixing of the full stained samples. First, clusters of cells with similar autofluorescence spectra are identified by unbiased dimensional reduction and clustering. Then, unique autofluorescence clusters are determined and are used to improve the unmixing accuracy of the full stained sample. This unbiased method allows for the identification of all autofluorescence spectra present in a sample, independent of cell types and intensity of the autofluorescence spectra. Furthermore, this method is equally useful for spectral analysis of different biological samples, including tissue cell suspensions, peripheral blood mononuclear cells and *in vitro* cultures of (primary) cells.

## Introduction

Autofluorescence is an intrinsic factor of cells that is caused by the natural emission of light by its cellular content and can complicate analysis methods in which fluorescence is used, such as imaging and flow cytometry (1–9). Different samples can have variable autofluorescence signatures depending on cell type, metabolic state, sample preparation and staining protocols, which can result in samples containing a heterogeneous autofluorescence profile (10). In addition, autofluorescence has been described as a reliable *in vitro* marker of cellular senescence and aging in several cell types, including human mesenchymal stromal cells (1), human nerve cells (11), human skin (12) and nematodes of C. elegans (13). As such, aged cells *in vitro* have increased autofluorescence.

Spectral flow cytometry is a powerful technology for cellular multiparameter analysis, which uses a mathematical algosrithm to define which photon corresponds to which unique spectral signature within a sample, using pure spectral signatures of fluorochromes as references, a process called spectral unmixing (14). Furthermore, this technology allows to fully characterize the unique autofluorescence profile within a sample, that is created by the different autofluorescence signatures of every cell within that sample (10).

In comparison to conventional flow cytometry, spectral flow cytometry captures the entire spectral profile of fluorochromes and autofluorescence by an array of detectors over multiple lasers that covers the entire range of emission wavelengths. In most spectral flow cytometry software packages, one autofluorescence signature per sample can be determined, allowing to distinguish between autofluorescence and fluorochrome signal (14, 15). However, when samples have a heterogeneous autofluorescence profile, it is challenging to resolve the real signal of all fluorochromes using the automated tools of the software, due to their limitations to deal with multiple autofluorescence spectra within one sample. A solution for this would be to add the different autofluorescence spectra as new references to the unmixing. Indeed, it has been shown that addition of one autofluorescence spectrum for a bright autofluorescence cell subtype, namely alveolar macrophages within in a murine lung cell suspension, was able to reduce the autofluorescence signal and improve the fluorochrome signals (16). For this, the cell type with the brightest autofluorescence had to be identified and gated based on FSC and SSC. However, when more cell types are present with overlapping FSC and SSC, and all with their own unique autofluorescence spectra, selecting autofluorescence cell types using these parameters will not be possible. The solution for this would be to identify every unique autofluorescence spectrum and add them all to the analysis.

A suggested workflow to identify unique autofluorescence subsets of cells would be to visualize unstained cells in a NxN matrix across all raw fluorescent detectors (10, 15). Sequen-tial gating of cells with different autofluorescence spectra in a bivariate manner and evaluation of their uniqueness would result in identifying some unique autofluorescence subsets. However, this laborious approach is subjective, requires a certain level of expertise and can only be used if the autofluorescence subsets are abundantly present and therefore can be visually detected (10). Moreover, autofluorescence subsets can easily be missed as the number of dimensions to check makes it complicated to capture all autofluorescence subsets manually. Therefore, this currently-used method is not sufficient for all types of samples, especially when containing many different autofluorescence spectra. Dimensional reduction of the unstained sample based on the signal intensity within the different detectors of a spectral flow cytometer can be used as an alternative approach (16). This will result in a bivariate plot, such as UMAP or tSNE, displaying heterogeneity based on intrinsic autofluorescence. Although this method helps to visualize different autofluorescence subsets, it still relies on manually selecting and gating the cells of interest.

Here, we provide an optimized approach to identify all different autofluorescence signatures within an unstained sample by including an unbiased clustering. Next, unique autofluorescence spectra are identified based on their similarity index and are subsequently used in the unmixing algorithm to extract the autofluorescence signal and improve the fluorochrome signal. We show that this approach is beneficial for various types of samples, including tissue cell suspensions with both heterogeneous and bright autofluorescence spectra (e.g. human lymph node cell suspension), cells grown *in vitro* with extremely bright autofluorescence spectra (e.g. human fibroblastic reticular cells (FRCs)) and also (primary) cell suspensions with heterogeneous and dim autofluorescence spectra (e.g. peripheral blood mononuclear cells (PBMCs)). Using this methodology, we identified that the unique autofluorescence spectra differ between donor samples and over time in one cell type upon culturing, highlighting the importance of following this workflow with every analysis for identification of the unique autofluorescence spectra in a sample in order to improve spectral data analysis and interpretation.

## Material and Methods

### Samples

Human lymph nodes were obtained from donors during liver transplant procedures performed at the Erasmus MC, Rotterdam, the Netherlands. The use of these tissues was approved by the medical ethical committee of the Erasmus MC (MEC-2014-060) and the liver transplant recipients provided written informed consent for the use of samples. Donor characteristics are included in Table S1. Lymph nodes were transported in Belzer UW cold storage solution (UW, Bridge to Life) on ice and processed within 72 hours after surgery. Lymph node cell suspensions were obtained through enzymatic tissue digestion as described previously (17). In short, lymph nodes were digested using RPMI-1640 supplemented with 2.4 mg/ml Dispase II, 0.6 mg/ml Collagenase P, and 0.3 mg/ml DNase I (all from Sigma-Aldrich) for 4 rounds of 10 minutes at 37 °C. After each round the isolated cells were washed in ice cold PBS supplemented with 2% FCS and 5 mM EDTA to stop the enzyme reaction and prevent overdigestion. Next, the cells were spun down at 300G for 4 minutes (4 °C), the cell pellet was resuspended in 1 ml DMEM with 10% FCS and passed through a 100 μm filter.

Peripheral blood mononuclear cells (PMBCs) were isolated from buffy coats obtained from healthy donors (Sanquin, The Netherlands) by density gradient centrifugation with Ficoll-Paque PLUS (GE Healthcare). No donor specific information is provided.

### Cell culture

Lymph node cell suspensions were plated at a minimum cell concentration of 20 x 10^6^ cells in a T-25 flask at 37 °C, 5% CO_2_. To grow out FRCs, flasks were pre-coated with 2 μg per cm^2^ using a solution of 50 μg/ml collagen from calf skin in HBSS (Sigma-Aldrich) and the cells were grown in DMEM supplemented with 10% FCS, 2% Penicillin/Streptomycin/Glutamine and 1% Insulin-Transferrin-Selenium (Gibco). After 3 days, lymphocytes were washed away with PBS in order to allow stromal cell growth. Once confluent, FRCs were harvested with 0.5% Trypsin + 5mM EDTA and passaged or used for flow cytometry.

### CD45^-^ stromal cell enrichment

Lymph nodes were digested as described above and enriched for CD45^-^ stromal cells by negative selection using MojoSort™ Human CD45 Nanobeads (Biolegend). CD45^-^ cell enrichment was performed according to manufacturer’s protocol, with the addition of a fluorochrome-labelled antibody against CD45 (Table 1) during the CD45 Nanobeads incubation step to allow optimal CD45 fluorescent signal for flow cytometry.

**Supplementary Note 1: Table 1.**
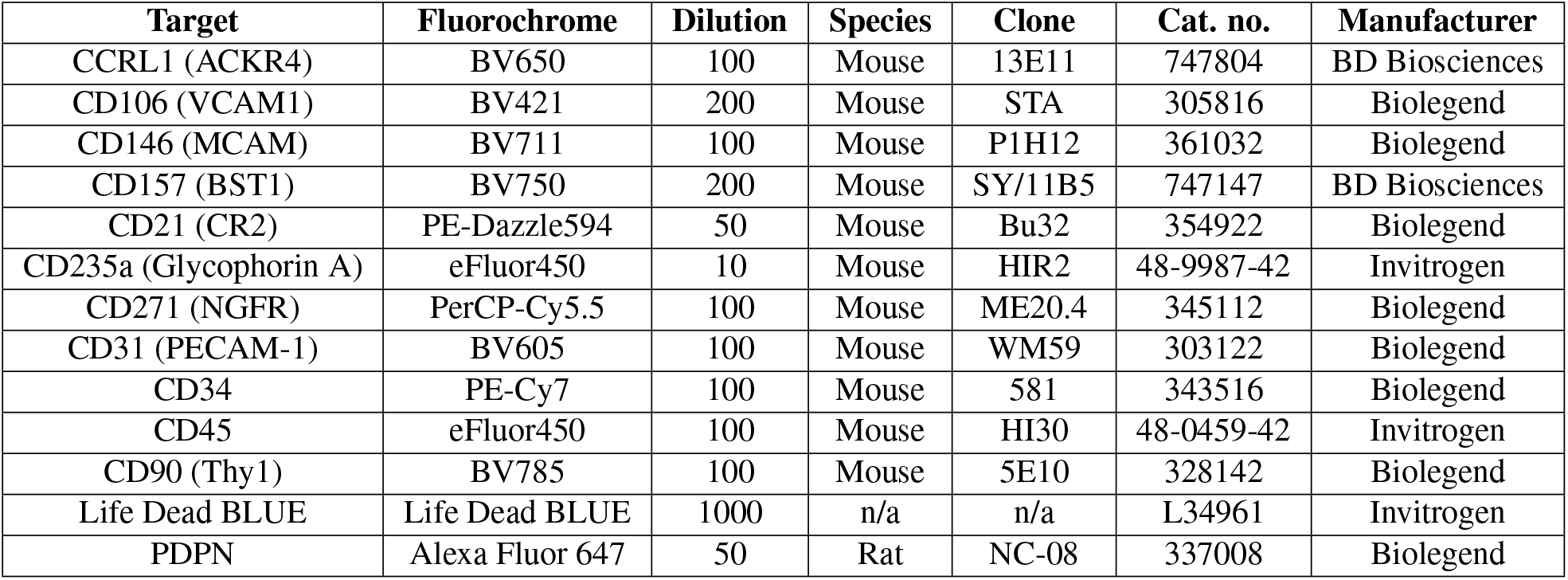
Antibodies used for flow cytometry.

### Flow Cytometry

Cell suspensions were stained in a 96-well U bottom plate on ice for flow cytometric analysis. Cells were firstly washed with FACS buffer containing PBS + 1% FCS, then washed with PBS and stained with a fixable viability dye (Table 1) for 10 min on ice. Fc-receptor blocking was performed using 10% normal human serum. The cells were then incubated with fluorochrome-labelled antibodies (Table 1). After staining, cells were fixed with 2 % PFA in PBS for 15 min, washed, and acquired on a 5 Laser Aurora spectral analyzer (Cytek Biosciences, Fremont CA). For appropriate autofluorescence spectra detection and unmixing, an unstained sample was taken along during the procedure, that was treated according to exactly the same procedure as the stained sample, of which at least the same amount of cells as in the stained sample were acquired. UltraComp eBeads compensation beads (ThermoFisher) were used as single stain controls for all fluorochrome-labelled antibodies, and cells heat shocked for 3 minutes at 65 °C were taken along as single stain control for the viability dye.

## Data analysis

All autofluorescence spectra in an unstained sample were visualized and identified by performing dimensional reduction (opt-SNE) and clustering (PhenoGraph) using the online software OMIQ (www.omiq.ai). The raw FCS file of the unstained sample was loaded into OMIQ and all detectors were scaled to 6000 Arcsinh, the default scaling setting in OMIQ. Opt-SNE and PhenoGraph were calculated based on all 64 detectors of the Aurora 5L and the identified PhenoGraph clusters were gated and exported as one FCS file. The R-script OMIQ-FCS-Separate was used to split this file into separate FCS files per cluster. These files were used in a new experiment in SpectroFlo^®^ to determine the similarity scores, by comparing the normalized spectra of all autofluorescence signatures. For this, a new fluorochrome library was created in SpectroFlo^®^ containing all different ‘autofluorescence signatures’, instead of fluorochromes. A reference group was assembled into a new experiment including an additional negative control and the ‘autofluorescence signatures’ corresponding with the number of autofluorescence clusters. An unstained control with an autofluorescence signal close to the background of the instrument was generated by acquiring 30% Contrad^®^ 70 (Decon). This unstained control was imported into the unstained and negative control tubes. The different autofluorescence clusters were imported as additional ‘autofluorescence signatures’ in the reference group, visually checked whether every cluster contained one pure spectra and used in the unmixing algorithm to define their uniqueness by means of calculating the similarity matrix. This matrix shows the similarity index (SI) of two spectra, which is a score between 0 and 1 that measures how closely two normalized spectra match. The unmixing algorithm of SpectroFlo^®^, that uses the Ordinary Least Squares (OLS), can unmix the sample when this number is ≤ 0.98 and there are at least 300 events per spectrum (15, 18). To identify the unique autofluorescence spectra, the similarity matrix was analyzed by the R-script SpectroFlo-Find-Unique-Spectra. Herein, the unique spectra are determined based on a ranking of the number of firstly, similar spectra (SI ≤ 0.98) and secondly, the sum of SI per cluster.

Next, the non-unique autofluorescence spectra were eliminated and the unique autofluorescence spectra were used to unmix the raw data of the original experiment, including unstained and full stained samples. The corresponding autofluorescence spectra were added to the reference group together with an additional negative control, to correct for the background noise of the machine. Again, the negative control and unstained sample of the reference group were replaced with a FCS file containing events without any autofluorescence and unmixing was performed.

The addition of extra fluorochromes to the unmixing can reduce the resolution of some fluorochromes, due to overlap between the newly added autofluorescence signatures and the spectra of the actual fluorochromes, leading to spillover spread. To improve the resolution, but still resolve the autofluorescence, the unmixing was evaluated after the reduction of the different autofluorescence spectra (Figure S1). The autofluoresence spectra were removed one by one from lowest to highest uniqueness, as given by the order of the output of the unique spectra identified with the R-script SpectroFlo-Find-Unique-Spectra, which is from most to least unique. After unmixing, the NxN plots of the unstained sample were evaluated to identify whether the removed autofluorescence spectrum was necessary to resolve all autofluorescence. If no autofluorescence was detected, this autofluorescence spectrum could be removed. However, if the autofluorescence was increased in the unstained sample, this autofluorescence spectrum could not be removed and was added back to the unmixing. Then, the next autofluorescence spectrum was removed from the unmixing and the whole procedure was repeated until all autofluorescence spectra were tested (Figure S1). After unmixing with the optimal number of autofluorescence spectra, live cells were gated and subsequently spillover correction was performed using SpectroFlo^®^.

Data is visualized using FCS express (De Novo Software) for the raw spectra and Prism (Graphpad Software) for the unique and normalized spectra and bar graphs. All other figures are produced using OMIQ (www.OMIQ.ai). For the density plots produced with OMIQ, debris was excluded and scatter plots (density) with 5 num levels and 0.1 percentile outliers was used to visualize the data.

## Data availability

The R-scripts used in this manuscript are publicly available at https://github.com/MolecularCellBiologyImmunology/Autofluorescence-Workflow.

## Results

### Determination of all unique autofluorescence subsets of cells using unbiased clustering

To correctly identify all autofluoresence spectra within a cell suspension for flow cytometric analysis, a workflow was created to identify all autofluorescence clusters present in a sample, with each cell cluster having its own unique autofluorescence spectrum (Figure 1A). First, the spectral profile of an unstained sample was acquired. To take into account autofluorescence induced during the preparation of the sample, the unstained sample had undergone the same wash-steps as the full stained sample. Furthermore, to be able to identify all different autofluorescence subsets, the same amount of events for the unstained sample as for the full stained samples was acquired.

**Fig. 1.**
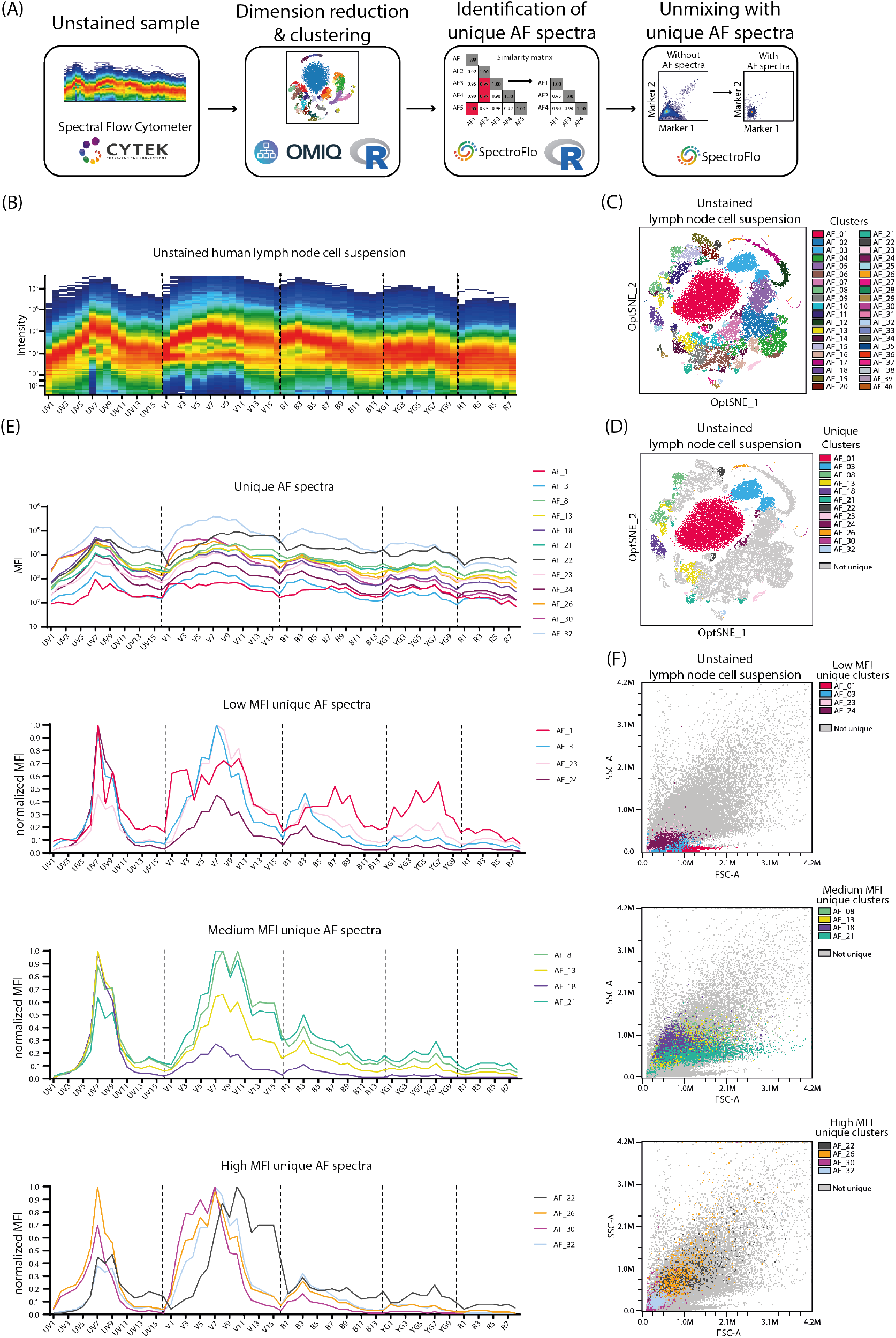
Identification of all unique autofluorescence spectra using dimension reduction and clustering. A) Schematic representation of the workflow and software needed to identify all unique autofluorescence spectra within an unstained sample that can be used for unmixing. B) Raw spectrum of an unstained human lymph node cell suspension acquired on Aurora 5L. C) Opt-SNE projection of all 40 autofluorescence clusters of the unstained human lymph node cell suspension. The Opt-SNE and PhenoGraph clusters were analyzed based on all 64 raw fluorescence detectors of the Aurora 5L. D) Opt-SNE projection of the 12 unique autofluorescence clusters of the unstained human lymph node cell suspension. The unique autofluorescence clusters were determined using the similarity index from SpectroFlo^®^ and the R-script ‘SpectroFlo-Find-Unique-Spectra’. E) Spectra and normalized spectra of the 12 unique autofluorescence clusters of the unstained human lymph node cell suspension. F) Scatter plots of the 12 unique autofluorescence clusters of the unstained human lymph node cell suspension. AF: autofluorescence, UV: ultraviolet, V: violet, B: blue, YG: yellow green, R: red, MFI: mean fluorescent intensity, FSC-A: forward scatter area, SSC-A: side scatter area.

To visualize and verify the different steps of the workflow that is presented in this paper, we used a freshly digested human lymph node cell suspension, as the lymph node contains different cell types including lymph node stromal cells, which tend to have highly heterogeneous and bright autofluorescence signatures. The signal intensities within the 64 detectors form a heterogeneous spectral profile in combination with high autofluorescence, with signals up to 10^6^ for the ultra-violet, violet, blue and yellow-green lasers (Figure 1B). The dimensional reduction opt-SNE and clustering algorithm PhenoGraph visualized and identified 40 clusters based on the signals from the 64 different detectors (Figure 1C). Next, it is important to identify which of these clusters are unique, i.e. have autofluorescence spectra that are different enough to be used by the unmixing algorithm. The similarity matrix, which is a measurement how closely spectra match, of all autofluorescence spectra is calculated in SpectroFlo^®^ and the custom made R-script identified the unique spectra based on both the ranking of the number of similar spectra (SI*≤* 0.98) and the sum of SI per cluster, as described before (10). For this human lymph node cell suspension, 12 unique autofluorescence clusters could be identified (Figure 1D). The unique autofluorescence clusters have autofluorescence spectral profiles with differences in brightness and with distinct normalized spectra (Figure 1E). Furthermore, the unique clusters could not be distinguished based on FSC and SSC (Figure 1F), showing that although the unique spectra identified are different from each other, they could not all have been identified based on the FSC and SSC parameters of the cells.

### Improved unmixing by addition of unique autofluorescence spectra

Next, the identified unique autofluorescence spectra were used for unmixing by adding them as separate autofluorescence signatures in addition to the fluorochromes already present in the antibody panel. Unmixing with additional 12 unique autofluorescence spectra reduced the autofluorescence signal present in the unstained and stained sample for different fluorochromes, including fluorochromes with peak emissions in the highly autofluorescence violet channels (BV605 & eFluor450), but also for fluorochromes with peak emissions overlapping with secondary emission channels from other fluorochromes (BV785 & PE-Cy7, BV711 & PerCP-Cy5.5) (Figure 2A-B). The CD45^+^CD31^+^ population, in which the hematopoietic marker CD45 is expressed by CD31^+^ cells, which are non-hematopoietic stromal cells, seen in the original unmixed sample, was incorrectly and no longer seen upon unmixing with 12 autofluorescence spectra (Figure 2B), showing that the signal of this population was created by autofluorescence and not by true fluorochrome signals.

**Fig. 2.**
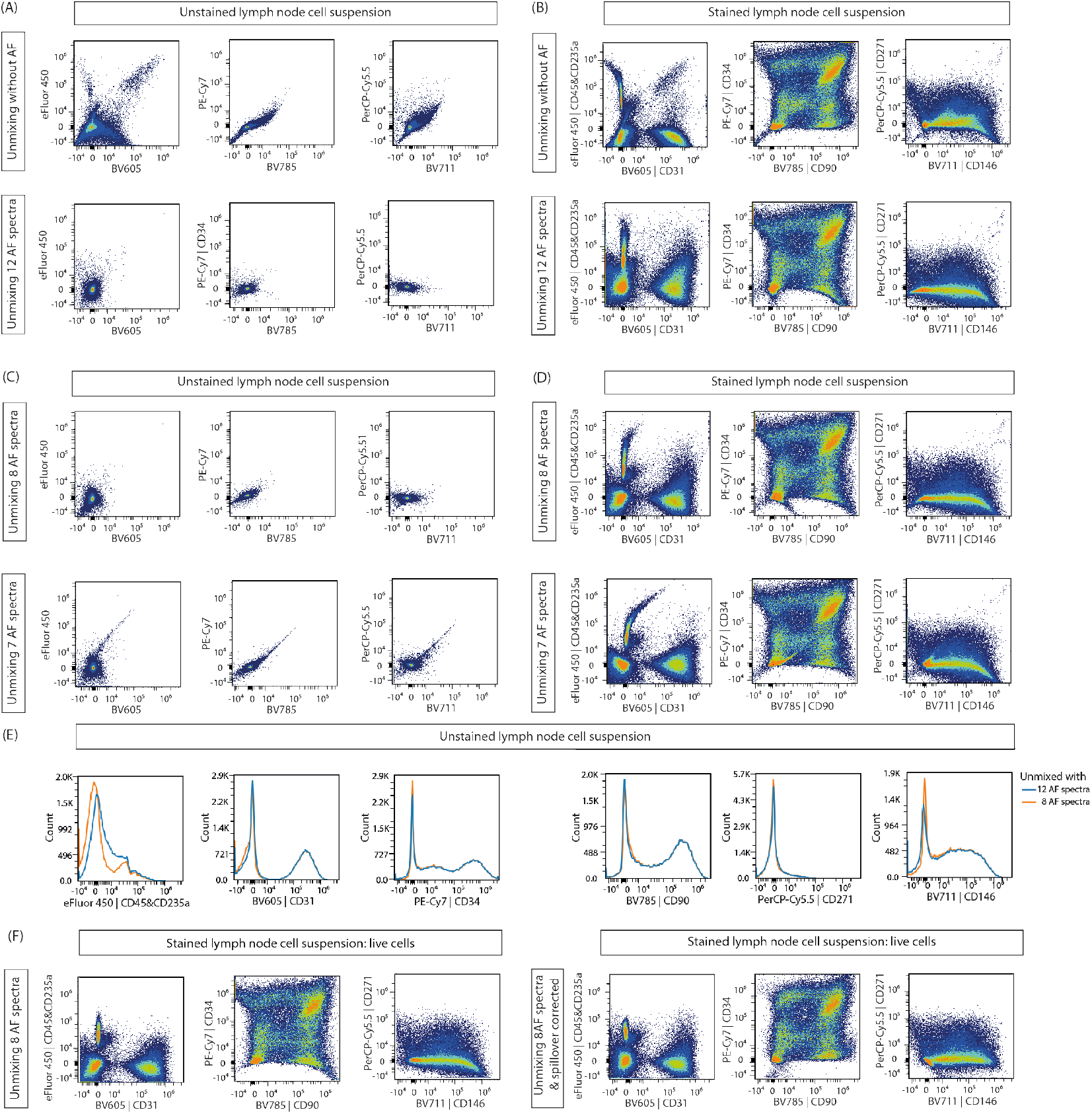
Unmixing of samples with or without the addition of autofluorescence spectra. A) Scatter plots of the unstained lymph node cell suspension after unmixing without (top) autofluorescence or with (bottom) the 12 unique autofluorescence spectra. B) Scatter plots of the stained lymph node cell suspension after unmixing without (top) autofluorescence or with (bottom) the 12 unique autofluorescence spectra. C) Scatter plots of the unstained lymph node cell suspension after unmixing with a reduced number of unique autofluorescence spectra, respectively 8 (top) and 7 (bottom) autofluorescence spectra. The spectra were reduced one by one from lowest to highest uniqueness (Figure S1). D) Scatter plots of the stained lymph node cell suspension after unmixing with a reduced number of unique autofluorescence spectra, respectively 8 (top) and 7 (bottom) autofluorescence spectra. E) Histograms of the stained lymph node cell suspension after unmixing with 12 (blue) or 8 (orange) unique autofluorescence spectra. F) Scatter plots of the stained lymph node cell suspension unmixed with 8 autofluorescence spectra, gated on live cells (left) and spillover corrected (right). For all plots, the fluorochromes visualized have peak emissions in the highly autofluorescence violet channels (BV605 and eF450), or have peak emissions overlapping with secondary emission channels from other fluorochromes (BV785 and PE-Cy7; BV711 and PerCP-Cy5.5). AF: autofluorescence, BV: brilliant violet.

The addition of extra autofluorescence spectra to the unmixing also resulted in lower resolution of the fluorochrome signal markers (Figure 2B). To improve the resolution of some markers while still resolving autofluorescence, we removed the autofluorescence spectra one by one from lowest to highest uniqueness (Figure S1). For this human lymph node cell suspension, the 12 unique autofluorescence spectra could be reduced to 8 unique autofluorescence spectra without increasing the autofluorescence signal (Figure 2C). In contrast, when the unmixing was reduced to 7 unique autofluorescence spectra, autofluorescence was again detected (Figure 2C). The reduction from 12 to 8 autofluorescence spectra during the unmixing especially improved the resolution of the fluorochrome eFluor 450 (Figure 2D-E). After unmixing with 8 autofluorescence spectra, the spread identified for some of the markers, especially PE-Cy7 vs BV785 and PerCP-Cy5.5 vs BV711, is reduced by spillover correction after gating on live cells (Figure 2F).

### Unbiased autofluorescence finding is beneficial for different types of samples

Identification of all unique autofluorescent spectra within a sample is beneficial for the unmixing of samples with both different cell populations, as well as cells with high autofluorescence, as seen for the human lymph node cell suspension (Figure 2). To test that this workflow is also valuable for spectral flow cytometry analysis of cells with high autofluorescence *in vitro*, and for heterogeneous cell suspensions with low autofluorescence, we analyzed fibroblastic reticular cells (FRCs) and PBMCs respectively. FRCs have extremely high autofluorescence, with signal up to 10^6^ in the ultraviolet, violet and blue channels and signal up to 10^5^ in the yellow-green and red channels (Figure 3A-i). Within this sample of FRCs, 6 different unique autofluorescence spectra were identified (Figure 3A-ii). Addition of these 6 autofluorescence spectra to the unmixing reduced the autofluorescence signal from unstained cells from 10^4^ to around 0 into different fluorochromes, including PE-Cy7 and BV785 (Figure 3A-iii). PMBCs have a heterogeneous spectral profile with autofluorescence signal up to 10^4^ in the ultraviolet, violet and blue channels (Figure 3B-i). Within this sample, 4 different unique autofluorescence spectra were identified (Figure 3B-ii). Addition of these 4 unique spectra to the unmixing reduced the autofluorescence signal in the unstained PMBCs in different fluorochromes, including eFluor450 and BV605 (Figure 3B-iii).

**Fig. 3.**
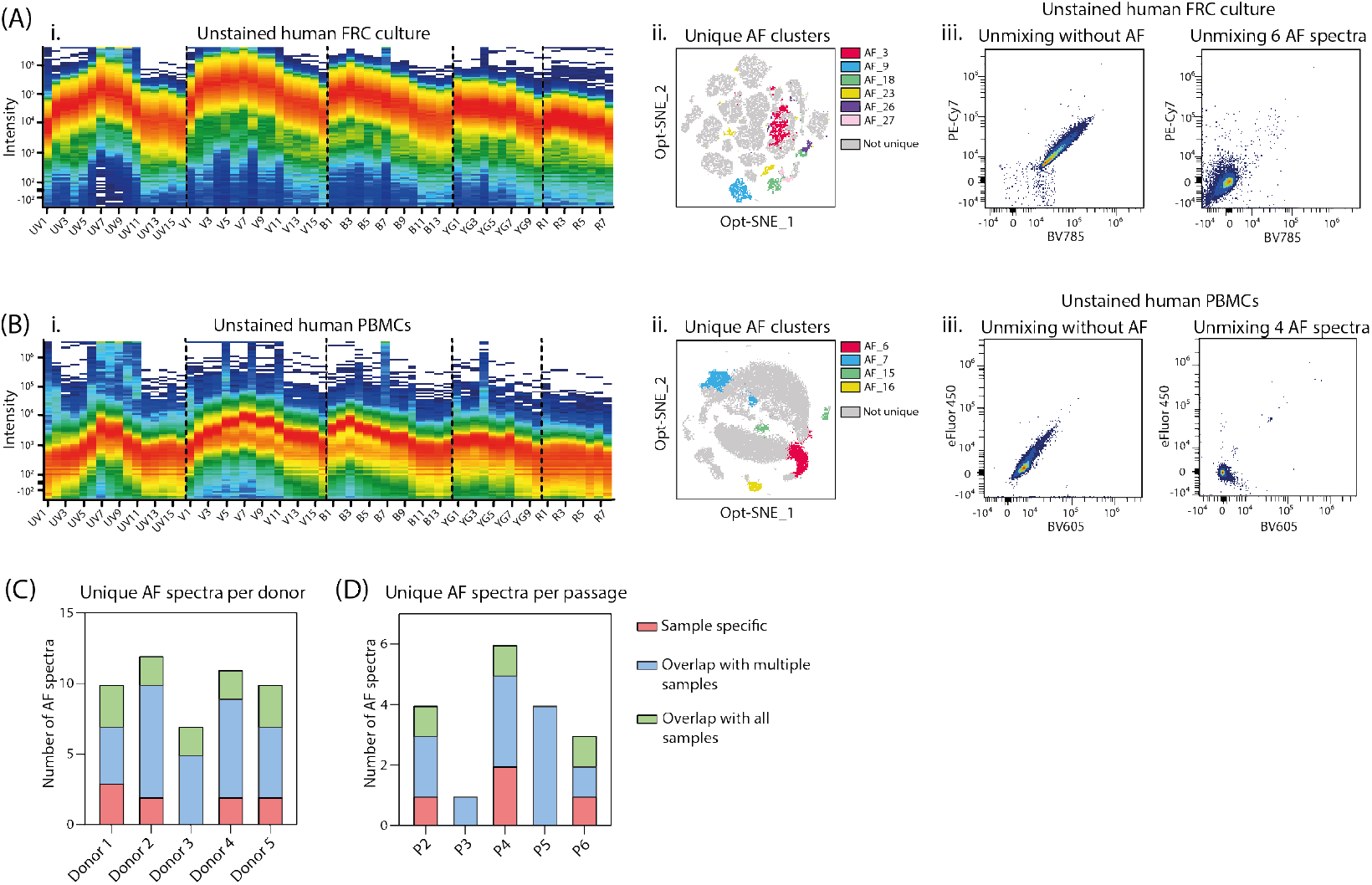
Unbiased autofluorescence detection in different types of samples. A) i. Raw spectrum of a sample of unstained human FRCs in culture acquired on Aurora 5L. ii. Opt-SNE projection of the 6 unique autofluorescence spectrum identified in this sample. iii. Scatter plots after unmixing without autofluorescence or with the 6 unique autofluorescence spectra. B) i. Raw spectrum of a sample of unstained human PMBCs acquired on Aurora 5L. ii. Opt-SNE projection of the 4 unique autofluorescence spectrum identified in this sample. iii. Scatter plots after unmixing without autofluorescence or with the 4 unique autofluorescence spectra. C) Count of the unique autofluorescence spectra in the human lymph node cell suspension of 5 different donors, divided into sample-specific (red), overlap with multiple samples (blue) or overlap with all samples (green). D) Count of the unique autofluorescence spectra per passage of FRCs from the same donor in culture for 5 passages, divided into sample-specific (red), overlap with multiple samples (blue) or overlap with all samples (green). FRC: fibroblastic reticular cell, UV: ultraviolet, V: violet, B: blue, YG: yellow green, R: red, AF: autofluorescence, BV: brilliant violet, PBMCs: Peripheral blood mononuclear cells, P: passage.

### Unique autofluorescence spectra differ between donor samples and within one cell type over time

The addition of one predefined autofluorescence spectrum per cell type have been shown to be beneficial for unmixing, for example the addition of the alveolar macrophage autofluorescence spectrum to lung cell suspensions (16). To identify whether this workflow can be used to create a library of unique autofluorescence spectra per sample or cell type that can be used for the unmixing of new samples of the same type, we compared the unique autofluorescence spectra found in different unstained samples. The unique autofluorescence spectra identified in lymph node cell suspensions of 5 different donors, which were processed and acquired on different days, did not completely overlap between the donor samples (Figure 3C). The observed differences were not related to the sex or the age of the donor (age 16 versus 83, Table S1). Although some unique spectra did overlap between all 5 donors, 4 out of 5 donors also had unique autofluorescence spectra that were specific for that donor sample (Figure 3C). A similar pattern is seen when comparing the unique autofluorescence spectra identified from passage 2 until 6 of cultured FRCs from the same donor. Three out of 5 passages had autofluorescence spectra overlapping with all other passages (Figure 3D). However, they also contained unique autofluorescence spectra that were not identified in other passages (Figure 3D). These data show that unique autofluorescence spectra differ between donor samples and within one cell type over time in culture and that selecting predefined autofluorescence spectra per cell type will not be sufficient to cover all autofluorescence within a sample. As such, this method needs to be performed prior to data analysis of each spectral flow cytometry experiment.

## Discussion

Here we introduce an unbiased workflow for spectral flow cytometry to identify all autofluorescence signatures in a variety of samples containing heterogeneous autofluorescence profiles. This method improves signal-to-noise ratio in spectral flow cytometry data, but is also valuable for autofluorescence-based cell sorting. Our method entails first the identification of all autofluorescence subsets using dimensional reduction (Opt-SNE) and clustering (PhenoGraph), then evaluation and selection of unique autofluorescence spectra based on the similarity matrix, after which the autofluorescence signal is extracted by adding them to the unmixing algorithm.

Recently, a workflow was published which also used dimensional reduction to visualize different autofluorescence subsets, however next cells with the highest autofluorescence spectrum were manually selected (16). We have now improved this workflow by replacing the manual gating with unbiased clustering and automated gating. The advantage of unbiased clustering is that this will select all subsets with unique autofluorescence spectra and not only the ones manually selected. Furthermore, some of the identified unique autofluorescence subsets within the lymph node cell suspension showed overlapping FSC and SSC profiles (Figure 1E-F). This suggests that, based on their autofluorescence profile, they are different subtypes with respect to autofluorescence even though they are similar with respect to FSC and SSC parameters. Thus, it is impossible to gate these different subtypes solely based on their FSC and SSC profile, additionally highlighting the need for unbiased clustering.

Besides the use of this workflow to improve signal-to-noise ratio, the identified unique autofluorescence signatures can also be used as a parameter on its own. It has been shown that distinct autofluorescence spectra can be used to identify different cell subsets, for example to distinguish hepatoblast-like cells and mature cholangiocytes within murine fetal livers (19). Moreover, autofluorescence can be used as parameter for label-free sorting on conventional cell sorters, for instance of senescent cells from mesenchymal stromal cell cultures (20) or neutrophils from peripheral blood human granulocytes (21). Combining high-end spectral cell sorters and the workflow presented in this paper could facilitate the sorting of different (unknown) cell subtypes purely based on their autofluorescence profile.

It has been shown that the addition of only one or two autofluorescence spectra to the unmixing algorithm could improve signal-to-noise ratios. For example, the addition of the autofluorescence spectrum of murine alveolar macrophages improved the unmixing of murine spleen, lung and skin cell suspensions (16). Also, the addition of both the autofluorescence spectra of lymphocytes and monocytes/neutrophils as separate spectra, improved the unmixing output of human PBMCs (22). In our study, we showed that unmixing accuracy could be improved by the addition of multiple autofluorescence spectra, demonstrating the importance of identifying all different unique autofluorescence spectra in a sample for the final analysis.

The workflow described in this paper is focused on the use of the Aurora spectral flow cytometer and the accompanied SpectroFlo^®^ software from Cytek Biosciences. However, with the addition of some small adjustments this protocol can be used for different types of spectral flow cytometers. To do so, it is necessary that the raw spectrum profile can be exported and that additional spectra can be added to the unmixing algorithm. As the Cytek Similarity Index, which is a metric developed by Cytek Biosciences, is exclusive for the SpectroFlo^®^ software, a matrix of Pearson r correlations could also be used for the identification of unique spectra when working with other software. In our hands (data not shown) and as described before (16), these two methods had an almost perfect linear correlation.

Some spectral flow cytometry software have incorporated a tool to take the autofluorescence spectrum of a sample into account during the unmixing. For example, the SpectroFlo^®^ software can treat the autofluorescence spectrum of the sample, defined by gating on the cells of interest based on FSC and SSC, as a separate parameter and extracts it from the fluorescence data, if desired (10). This can be used when the sample contains a single autofluorescence spectrum, as this spectrum is extracted from all cells within the sample. However, this tool is not sufficient to reduce all autofluorescence in samples with heterogeneous autofluorescence profiles. As this tool extracts the mean autofluorescence spectrum from a mixture of cells with all different autofluorescence profiles, it would result in photons from autofluorescence being wrongly assigned. This would impact the resolution between the fluorochromes, making the analysis less sensitive. Therefore, in a heterogeneous sample it is necessary to identify all different autofluorescence subsets to use them in the unmixing algorithm.

The method described here is unbiased in identifying and selecting unique autofluorescence spectra. The only manual component described in our workflow is the reduction of the number of added autofluorescence spectra in the unmixing to improve the resolution of some fluorochromes. To make this workflow totally unbiased, a tool has to be created that can evaluate the effectiveness of the different combinations of autofluorescence spectra used for unmixing. For example, by generating an unmixing score based on the signals for all the different fluorochromes within the unstained sample, especially by paying attention to the outliers and extreme negative values. This can then be used to evaluate in an unbiased setting whether the reduction of added autofluorescence spectra affects the unmixing accuracy.

Remarkably, we showed that unique autofluorescence spectra are present in similar tissue samples from different donors and even within cultures of the same cells over different passages. Therefore, we highlight that this method should be performed with every new acquired sample, to identify the unique autofluorescence spectra for that specific sample at that specific time. However, when the same sample types are used, an additional step can be added to the workflow to build an extensive library of all unique spectra identified that could not be removed and were thus necessary for correct unmixing accuracy. When enough samples are acquired, these spectra can be evaluated to identify whether there is a pattern of autofluorescence spectra that are always needed for correct unmixing. These spectra can then be beneficial for the unmixing of later experiments with the same sample types. Although such a pattern was not present for the lymph node cell suspensions and *in vitro* FRCs (Figure 3C-D), we hypothesize that such a pattern might be present in other samples, depending on the sample type, number of samples, preparation and acquisition.

Besides the mathematical extraction of the autofluorescence signal after acquisition, reduction of the autofluorescence signal before acquisition is often used in fluorescent microscopy (23–25). Various reagents have been described to quench autofluorescence signal. For example Sudan Black B has been shown to eliminate autofluorescence on both frozen and paraffin embedded sections of multiple tissues (23, 24). We have explored whether these methods can also be applicable for flow cytometric analysis. Unfortunately, we noticed that Sudan Black B adds its own fluorescent spectrum to our flow cytometry samples, making it not useful for quenching autofluorescence in spectral flow cytometry (unpublished observations). Another method that has been described to reduce autofluorescence in microscopy is photobleaching using UV light, which induces irreversible modifications of fluorochromes (25). This method has also been described to quench antibody-conjugated fluorochromes signal on live cells in suspension. To minimize viability loss, constant cooling and anti-oxidants are necessary to prevent reactive oxygen species from damaging cells (26). Antibody-conjugated fluorochromes can be photobleached in 3 to 25 minutes. However, to quench autofluorescence a longer exposure time is needed: 24 hours to multiple days have been described for microscopy (25). Because a long exposure time to UV light would kill live, unfixed cells and quench fluorescent signals from antibody-conjugated fluorochromes, autofluorescence photobleaching could only be used for flow cytometry on fixed, unstained samples. This reduces its capacity as a general method to reduce autofluorescence in spectral flow cytometry. Overall, reduction of the autofluorescence before acquisition is not always feasible, and a simple way to get the best signal-to-noise ratio when working with autofluorescence samples is to assign the fluorochromes used within the antibody panel on forehand in such a way that they have the least overlap and interference with the autofluorescence spectrum of the unstained cells.

To summarize, we here describe an improved, unbiased method to identify and select all unique autofluorescence spectra using spectral flow cytometry. Addition of these spectra to the unmixing algorithm reduces the autofluorescence background and increases the resolution to detect different fluorochromes. This workflow is beneficial for spectral flow cytometry analysis of various types of samples, including samples containing very high autofluorescence signal and/or with a heterogeneous autofluorescence profile.

## ACKNOWLEDGEMENTS

The authors would like to thank and acknowledge the expert help of the Microscopy and Cytometry Core Facility at Amsterdam UMC location Vrije Universiteit, Amsterdam. We thank Kim Ober and Dr. Monique Verstegen from the Erasmus Medical Centre, Rotterdam, for sample logistics, and Dr. Maria Jaimes from Cytek for the critical reading of the manuscript.

## Author contributions

Conceptualization: **JEGR, JJGV, CMdW** and **REM**; Methodology: **JEGR, AMM, CHC, JJGV, CMdW** and **REM**; Investigation: **JEGR** and **AMM**; Software: **JEGR** and **MdK**; Data Curation: **JEGR** and **AMM**; Writing – Original draft preparation: **JEGR**; Writing – review and editing: **JEGR, JJGV, HPR, LJWvdL, CMdW** and **REM**; Supervision: **CMdW** and **REM**; Funding acquisition: **REM**; Ethical approval and donor information: **HPR** and **LJWvdL**. All authors have read and agreed to the published version of the manuscript.

## Ethical statement

Human lymph nodes were obtained from donors during liver transplant procedures performed at the Erasmus Medical Centre in Rotterdam, the Netherlands. The use of the tissues was approved by the medical ethical committee of the Erasmus MC (MEC-2014-060) and the liver transplant recipients provided written informed consent for the use of samples.

---

**Supplementary Note 2: Table S1.** Lymph node donor information. DBD: Donation after Brain Death; DCD: Donation after Circulatory Death; FRCs: fibroblastic reticular cells; LN: lymph node.

| Donor | Age | Sex | Donor type | Tissue/cell type used |
| --- | --- | --- | --- | --- |
| 1 | 83 | Female | DBD | LN cell suspension |
| 2 | 16 | Female | DBD | LN cell suspension |
| 3 | 21 | Female | DBD | LN cell suspension |
| 4 | 53 | Female | DBD | LN cell suspension |
| 5 | 65 | Male | DCD | LN cell suspension |
| 6 | 33 | Female | DCD | *In vitro* culture of LN FRCs |

**Supplementary Note 3: Figure S1.**
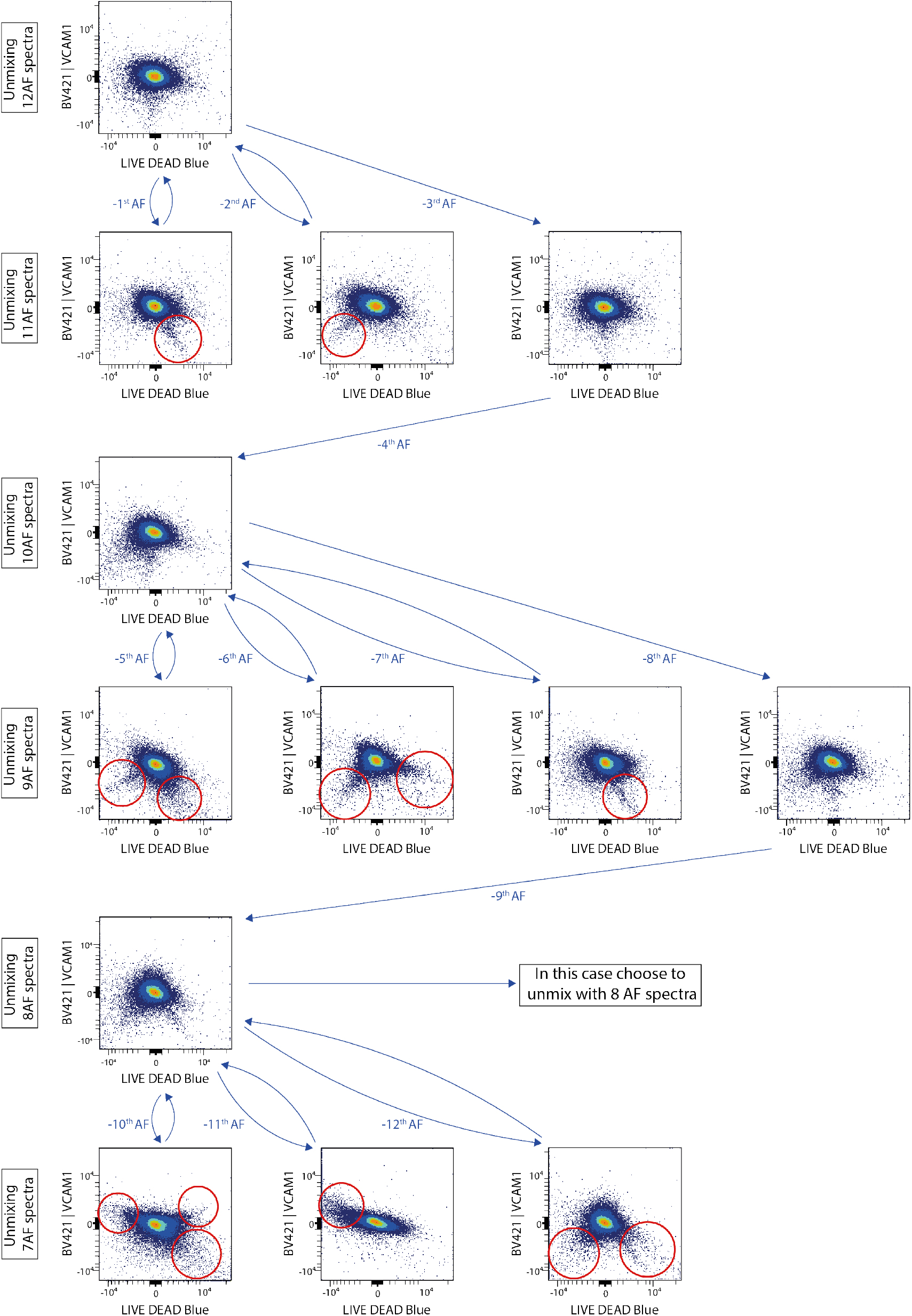
One-by-one reduction of the autofluorescence spectra during unmixing to identify the spectra necessary for unmixing. Visualization of the decision tree for the one-by-one reduction of autofluorescence spectra of the unstained human lymph node cell suspension. The autofluorescence spectra were removed one-by-one from lowest to highest uniqueness, after which the NxN plots of the unstained sample were evaluated to identify autofluorescence signal. The scatter plots of LIVE DEAD Blue and BV421 are shown as example. After removing the first autofluorescence spectrum, new autofluorescence signal was detected (red circle). Therefore, this spectrum could not have been removed and was added back. Then the second autofluorescence spectrum was removed, and again autofluorescence signal was detected (red circle). Therefore, this spectrum could not have been removed and was added back. Next the third autofluorescence spectrum was removed, which does not induce extra autofluorescence signal, so this spectrum could be removed. Afterwards, the fourth autofluorescence spectrum was removed and evaluated. This process was continued until all spectra have been removed and the autofluorescence signal was evaluated. For this sample we decided to continue the unmixing with 8 unique autofluorescence spectra. AF: autofluorescence, BV: brilliant violet.

